# A smartphone microscope method for simultaneous detection of (oo)cyst of *Cryptosporodium* and *Giardia*

**DOI:** 10.1101/2020.04.09.035147

**Authors:** Retina Shrestha, Rojina Duwal, Sajeev Wagle, Samiksha Pokhrel, Basant Giri, Bhanu Bhakta Neupane

**Author notes:** Corresponding email: BBN, BG.

## Abstract

Gastrointestinal disorders caused by ingestion of (oo)cysts of *Cryptosporodium* and *Giardia* is one of the major health problems in developing countries. We developed a smartphone based microscopic assay method to screen (oo)cysts of *Cryptosporodium* and *Giardia* contamination in vegetable and water samples. We used sapphire ball lens as the major imaging element to modify a smartphone as a microscope. Imaging parameters such as field of view and magnification, and image contrast under different staining and illumination conditions were measured. The smartphone microscope method consisting of ball lens of 1 mm diameter, white LED as illumination source and Lugols’s iodine staining provided magnification and contrast capable of distinguishing (oo)cysts of *Crypstopsporodium* and *Giardia* in the same sample. The analytical performance of the method was tested by spike recovery experiments. The spiking recovery experiments performed on cabbage, carrot, cucumber, radish, tomatoes, and water resulted 26.8±10.3, 40.1±8.5, 44.4±7.3, 47.6±11.3, 49.2 ±10.9, and 30.2±7.9% recovery for *Cryptosporodium*, respectively and 10.2±4.0, 14.1±7.3, 24.2±12.1, 23.2±13.7, 17.1±13.9, and 37.6±2.4 % recovery for *Giardia*, respectively. These recovery results were found to be similar when compared with the commercial brightfield and fluorescence microscopes. We tested the smartphone microscope system for detecting (oo)cysts on 7 types of vegetable (n=196) and river water (n=18) samples. Forty two percent vegetable and thirty-nine percent water samples were found to be contaminated with *Cryptosporodium oocyst*. Similarly, thirty one percent vegetable and thirty three percent water samples were contaminated with *Giardia cyst*. This study showed that the developed method can be a cheaper alternative for simultaneous detection of (oo)cysts in vegetable and water samples.

## Introduction

Food and waterborne illnesses arising from the consumption of contaminated food and water are serious health hazards globally [1]. The World Health Organization (WHO) has reported 1.5 billion episodes of diarrhoeal cases leading to 3.5 million deaths of under 5-year-old children in developing countries annually. More than 70% of these diarrhoeal episodes are attributable to biologically contaminated food [2]. In order to prevent and identify the disease, detection of food borne parasite is important at all levels of production chain followed by screening and certification [3].

*Cryptosporidium* and *Giardia* are the major food and water-borne parasites [4]. Ninety percent of reported outbreaks of these pathogenic protozoans occur through water, while 10% are related to food. In the infective stage, *Cryptosporidium* oocysts have spherical shape with a diameter of 4-6 μm and *Giardia* cysts have elliptical shape of 8–12 um long and 7–10 μm wide [5]. Both of the cysts, collectively termed as (oo)cysts, have a simple and direct life cycle, which is extremely suitable for transmission by fresh produce. Additionally, the cysts are small in size with a robust transmission stage. Some genotypes of the parasites even have zoonotic potential giving the opportunity for contamination to occur from both animal and human sources. Cryptosporidia are particularly threatening as they are resistant to chlorine disinfection, can persist in the environment for a long period, can infect other animal hosts, and are difficult to diagnose and treat. The infectious dose for *Giardia* cysts and *Cryptosporidium* oocysts are 10–100 and 10–1000 respectively, which makes these pathogens more precarious [6]. Developing countries are the most vulnerable countries to these protozoans where infection is more likely underdiagnosed and underreported, and has limited resources and infrastructures for investigation [7]. In low income countries, the overall prevalence rate of Giardia infection is 20 – 30% and the occurrence of *Cryptosporidium* are 4–31% in children younger than 10 years [8].

Several methods have been described to detect *Giardia* cyst and *Cryptosporidium* oocyst in food, water, and faecal samples with high sensitivity and specificity. Commonly used approaches are polymerase chain reaction, flow cytometry, optical microscopic examination etc. However, these techniques need a good laboratory facility, well trained user and are expensive, therefore are not appropriate for low–resource settings including remote and field sites. There is a need for a simple, easy to use, rapid but reliable and low–cost test method for the detection of parasites [9–12].

In recent years, smartphone based systems are being explored and used as an alternative platform for the detection of microscopic to sub–microscopic specimens in a wide variety of matrices, such as parasite eggs in faecal sample[13], allergen in food [14], blood cells in blood [15], single nanoparticles and viruses [16], filarial and malarial parasites in blood [17, 18], sickle cell anaemia in a blood smear [19], soil–transmitted helminth and fluke in urine and stool samples [20] etc.

In this work, we describe a smartphone microscopic system that can image and quantify (oo)cysts of both *Cryptosporidium* and *Giardia* in a given sample. We optimized optical parameters of the microscope including field of view, magnification, and image contrast under different staining and illumination conditions. The validity of the developed microscope was tested by spiking the vegetable sample with known number of standard (oo) cyst samples. For comparison, the spiked vegetable samples were also imaged with a commercial bright field and a fluorescence microscope and the percentage recovery data were compared. The optimized smartphone microscope was also used to measure (oo)cyst contamination in vegetable, fruits and water samples.

## Materials and methodology

### Design of smartphone microscope

We used a sapphire ball lens (Edmund Optics, New Jersey, USA) and a mounting plate made up of aluminium to transform a smartphone (Samsung Galaxy J7 prime) into a smartphone microscope. A small hole was punctured at the centre of the mounting plate and the ball lens was firmly glued in this hole (figure 1A). The ball lens was then centred over the smartphone camera lens. The mounting plate was fixed onto the smartphone using a transparent tape. The smartphone had a rare camera of 13 MP and the screen size of the phone was 151.7 mm x 75 mm with 1080 x 1920 pixels resolution. We custom built a microscope stage to hold the sample slide using a wooden viewing box of dimension 15×15 cm. A 3 cm diameter hole was drilled on top centre position of the box to have the illumination light pass through the sample specimen placed on the microscopic slide just above the hole. The slide was fixed on both side of the hole using a double-sided tape. A schematic of the optical set up is shown in figure 1.

**Figure 1:**
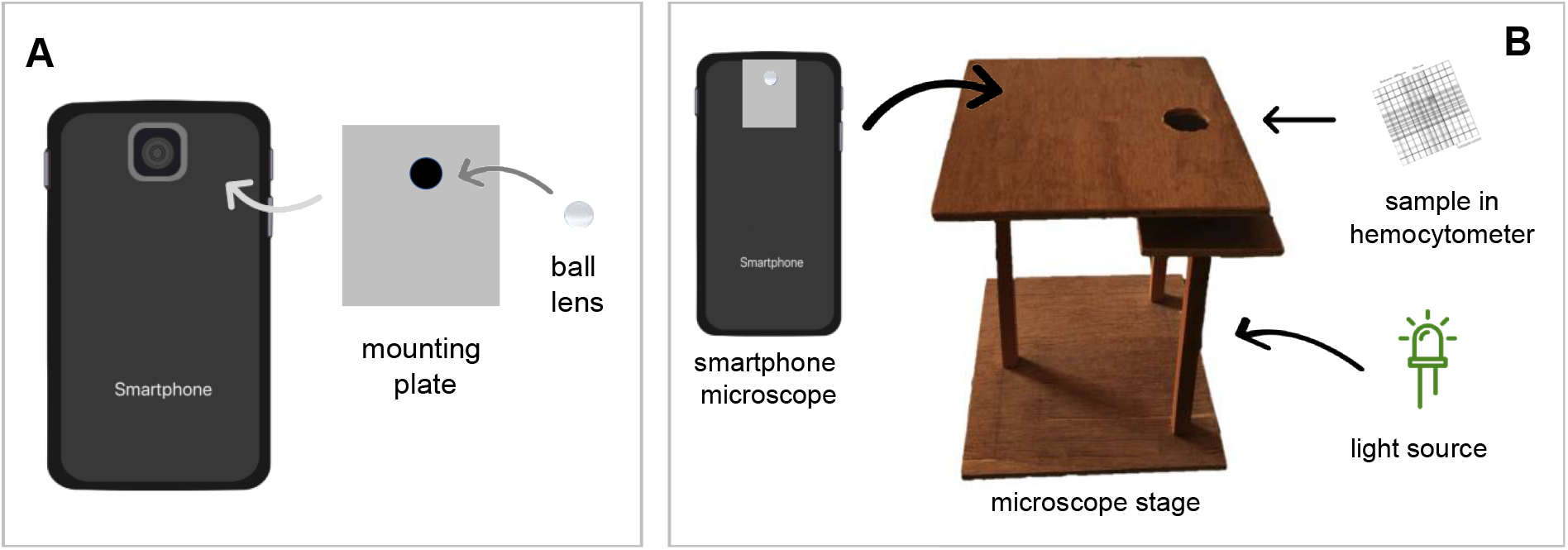
Assembly of smartphone microscope (A) and a measurement set up (B).

### Measurement of magnification and contrast

We tested three ball lenses having different diameter of 0.5 mm, 1 mm, and 2 mm. Image of a standard calibration grid (Generic, USA; 1 division = 10μm) was captured with the ball lens attached to the smartphone. The distance covered by all the grid lines was measured in pixel using ImageJ software. The pixel distance was converted to micrometre to get field of view (FOV) of the microscopic system.

The magnification (*MAG*) was calculated as:

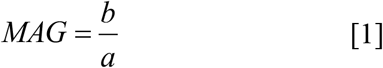

Where, *a* and *b* are size of object and image, respectively.

To measure the contrast, we imaged standard (oo)cysts sample (Waterborne TM, Inc., New Orleans, USA) under different illumination and staining conditions to determine the image contrast. Based on field of view and magnification, we chose 1 mm ball lens in our further experiments. We tested two different light sources for sample illumination: a smartphone flashlight and a white light emitting diode (LED, 12 watt). We also tested two different types of stains: Lugol’s iodine (HiMedia, Mumbai, India) and methylene blue dye (Fisher Scientific, New Hampshire, United States). The staining experiments involved a mixture of *Cryptosporodium parvum* and *Giardia intestinalis* (oo) cysts suspension (~ 2×10^5^ (oo) cyst per mL) and stain dye in 1:1 ratio. An aliquot of 10 μL of the mixture was loaded on the Haemocytometer (Max Levy, Philadelphia, USA) and was then illuminated by the light source and viewed through smartphone microscope. For each light source, images of (oo)cysts were captured at different time intervals after mixing the dye with the standard (oo)cysts. The images were analysed in ImageJ software to calculate the Weber contrast in percentage (WC) as follows [21]:

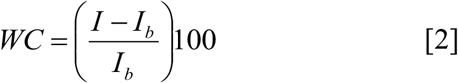

where, *I* is the intensity on the specimen of interest i.e. (oo) cyst and *I_b_* is the average intensity of immediately adjacent background.

### Spike recovery experiments

Five different types of vegetables such as tomato, cabbage, carrot, radish, and cucumber were selected for spike recovery experiments. These vegetables were selected based on previous reports of faecal contamination [22–25]. These vegetables are consumed in raw forms in many countries. All the samples were bought from the local vegetable shops. A portion of the sample (15-20 g) was soaked in distilled water for about 20 minutes to remove all the surface contaminants including (oo)cysts. After washing the samples, 10 μL of a mixture of *C. parvum* oocysts and *G. intestinalis* cyst suspension (Waterborne Inc, PC101 G/C positive control) was added to randomly selected points on sample surface samples with a micropipette. The (oo)cysts seeded sample was left to dry at room temperature for 2 hours to overnight.

We also performed similar spike recovery experiments with water samples. Five sets of 50 mL distilled water (pH= 6.8, conductivity= 0.01Sm^−1^) were spiked with 10 μL of the standard (oo)cyst suspension and were incubated overnight.

We tested three different washing solutions such as distilled water, normal saline, and glycine buffer (1M, pH 5.5) to extract (oo)cysts from each spiked vegetable samples The sample was put in extracting solution in a ziplock bag (Great Value, Fresh Seal Double Zipper). Ziplock bag was used as a low–cost alternative to stomacher bag. The eluate was then carefully transferred into two 50 mL falcon tubes. The elute suspension was concentrated by using the triple centrifugation method proposed by Medeiros and Daniel [26] with some modifications. At first eighty millilitres of eluate was centrifuged in two 50 mL tubes at 1500 x g for 10 minutes. The supernatant was decanted into a clean beaker leaving a final volume of 5 mL, which was placed in a vortex mixture for 20 seconds to homogenize the pellet. The 5 mL residual volume from each centrifugation tubes were combined together into a single tube. Another centrifugation was carried out at 1500 x g for 10 minutes. The supernatant was discarded leaving 0.5 mL pellet in the centrifuge tube. The residual solution was again vortexed for 20 seconds and it was carefully transferred to 1.5 mL microcentrifuge tube with 10 μL micropipette. The centrifuge tube was rinsed with 0.5 mL distilled water and added to the same 1.5 mL microcentrifuge tube to make the final volume of 1 mL. Now, the third centrifugation was performed at 1500 x g for 10 minutes. The supernatant was removed, leaving just 0.5 mL in the microcentrifugation tube one more time.

The water samples containing (oo)cysts were subjected to flocculation and sedimentation as described by Karanis and Kimura [27] with some modifications. 50 mL of ferric sulphate (0.25 M) solution was added to 50 mL of water samples and the pH was adjusted to 6±0.05. The sample was left 24 hours at room temperature to precipitate floc. Then the supernatant was carefully aspirated with a syringe filter without disturbing the sediment. The sediment was further centrifuged at 2,000 ×*g* for 10 minutes and the supernatant was discarded. The pellet was dissolved in 1 mL of citric acid lysis buffer (8.4 g citric acid monohydrate, 17.64 g tri–sodium–citrate–dihydrate, distilled H_2_O up to 100 mL; pH 4.7) by incubating at room temperature for 1 hour with vortexing every 15 minutes. The sample was washed twice with distilled water by centrifugation at 2000 ×*g* for 10 minutes. The pellets collected were resuspended in 5 mL distilled water for the purification of (oo)cyts. The purification step is only required with contaminated water, whereas non–contaminated water can be pelleted followed by dissolving in the buffer and subjected to the microscopy.

The purification steps of water samples involved a discontinuous sucrose gradient. The gradient was prepared with the Sheather’s solution (320 mL H_2_O and 500 g sucrose) diluted with 0.025 M phosphate-buffered saline (PBS) and supplemented with 1% Tween 80 to make 1:2 solution of 1.103 specific gravity and 1:4 solution of 1.064 specific gravity. 10 mL of 1:4 solution was layered over 10 mL of 1:2 solution on a 50 mL centrifuge tube. Then, 5 mL of the sample were layered over 1:4 solution and was centrifuged at 1500 ×*g* for 30 minutes. The three layers were recovered carefully and pooled separately along with the pellet and examined for the oocysts. The pooled layers were diluted with water, centrifuged and pellets were collected for the microscopic analysis.

### Microscopic measurements

Ten microliters of each concentrated sample were stained with 10 μL of diluted Lugol’s iodine (1:2 in water) and subsequently loaded into hemocytometer. The sample was incubated for around 6 minutes. The (oo)cysts were screened and enumerated in four quadrants of the hemocytometer under smartphone microscope. The cysts on the same hemocytometer were simultaneously counted by brightfield microscope. Triplicate measurement was made for each concentrated suspension.

The spiked samples were also examined with a fluorescent microscope (Labomed Inc, United States, LB 702). For fluorescence measurement, 5μL of (oo)cyst suspension was placed on the clean glass slide to which a drop of fluorescein–labelled mouse monoclonal antibodies (Aqua-Glo™ G/C) was applied. The slide was incubated at 37°C for 10 minutes in an incubator (Faithful, China) and imaged with 480 nm excitation and 520 nm emission wavelengths.

The numbers of (oo)cyst were counted before each seeded experiment using all three microscopic methods. The mean percentage recovery efficiency (RE) was calculated as:

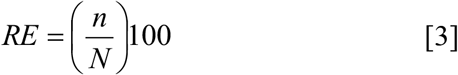

Where, *N* is the number of (oo) cysts added to a sample and *n* is the number of (oo) cysts recovered from the sample.

We followed the above procedure excluding the washing of raw vegetables and spiking with (oo)cyst to determine the (oo) cyst contamination in vegetable and water samples.

## Results and discussion

### Optimization of smartphone microscope

The performance of an imaging system is determined by its optical parameters, such as field of view, magnification, resolution, and contrast. The FOV is the size of the viewing area that can be sees when we look through a microscope. Magnification measures the zooming of an object, and resolution and contrast measure the details and clarity in an image [28]. The FOV of ball lens based imaging system depends on the size of ball lens, the refractive index of ball lens material and wavelength of illumination source; the size factor being major contributor [29]. On the other hand, the magnification of smartphone microscope depends on size of ball lens and also on the nature of smartphone that contains a built-in lens and CMOS camera at fixed distance. The spherical ball lens has a curved surface that results in curvature effect. It means the central region is more sharp/clear than the periphery in the image plane. The clear field of view microscope having ball lens of 0.5, 1, and 2 mm ball lens is provided in table 1.

**Table 1:** Field of view (FOV) of the smartphone microscope

| Diameter of ball lens<br>(mm) | Clear field of view<br>(μm) |
| --- | --- |
| 0.5 | 114±5.6 |
| 1 | 203±5.7 |
| 2 | 638.7±15.9 |

The measured field of view (FOV) of smartphone microscope showed that the FOV increases with an increase in the diameter of ball lens (Table 1) which is in consistent with reported values [29].

The magnification of smartphone microscope with 1 mm ball lens was estimated to be 200×. The 1 mm lens set up was able to magnify *Cryptosporodium* oocyst to 0.8–1.2 mm and *Giardia* cyst to 1.4–2.4 mm, respectively. The magnification was enough to distinguish the two specimens. Therefore, we selected a smartphone microscopic system having 1 mm diameter ball lens for further experiments in this study.

Light source used for illuminating sample affects the image quality of (oo) cyst. We tested following two types of commonly available light sources: white LED and smartphone light. The images collected using these light sources are shown in figure 2.

**Figure 2:**
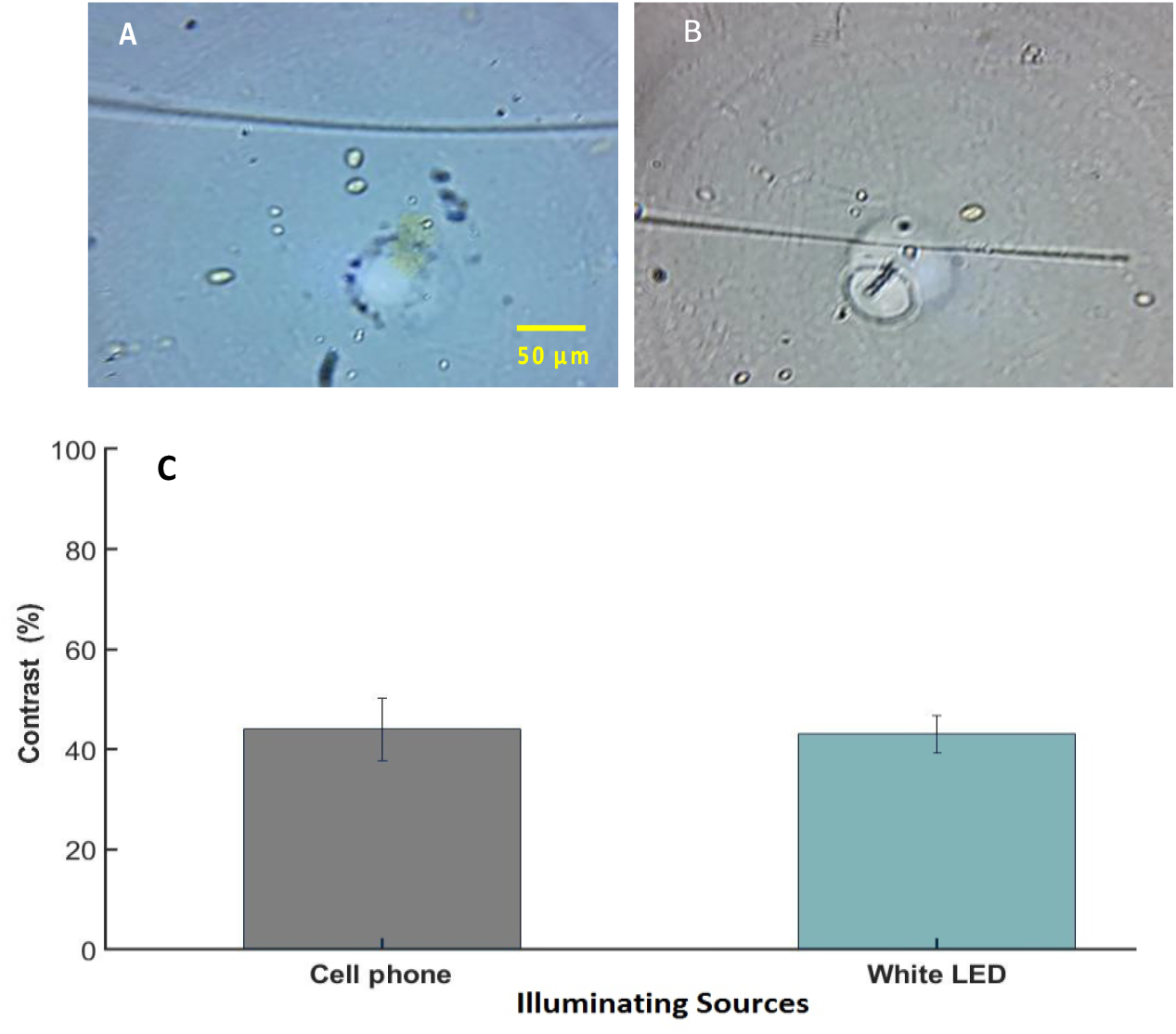
Photographs of (oo)cysts of cryptosporidium and giardia taken using smartphone microscope with 1 mm ball lens with (A) white LED light illumination and (B) smartphone flashlight illumination. Scale bar in both images is 50 μm. (C) The measured contrast for (oo)cysts. The error bar in C represent the standard deviation of triplicate measurements.

Images shown in figure 2 clearly depicts that the oocyst and cyst can be easily distinguished from each other based on their shape and size. Oocysts are circular shape and cysts are oval in shape. The Weber contrast for the (oo)cysts are shown in figure 2C. Both of the light sources provided similar contrast percentages. Since the white LED is easily available, cheaper, and easy to use, we chose it for further experiments.

A number of staining procedures have been developed to aid in the clear morphological identification and differentiation of (oo)cysts by light microscopy [9]. Some of the most used techniques are the iodine and methylene blue mounts. These methods are simple, faster and inexpensive and provide clear distinction of (oo)cyst by morphological features [10]. The temporal variation of stain color intensity on the cysts are shown in figure 3. This shows the color intake by the cysts and stability of the stains with waiting time.

**Figure 3:**
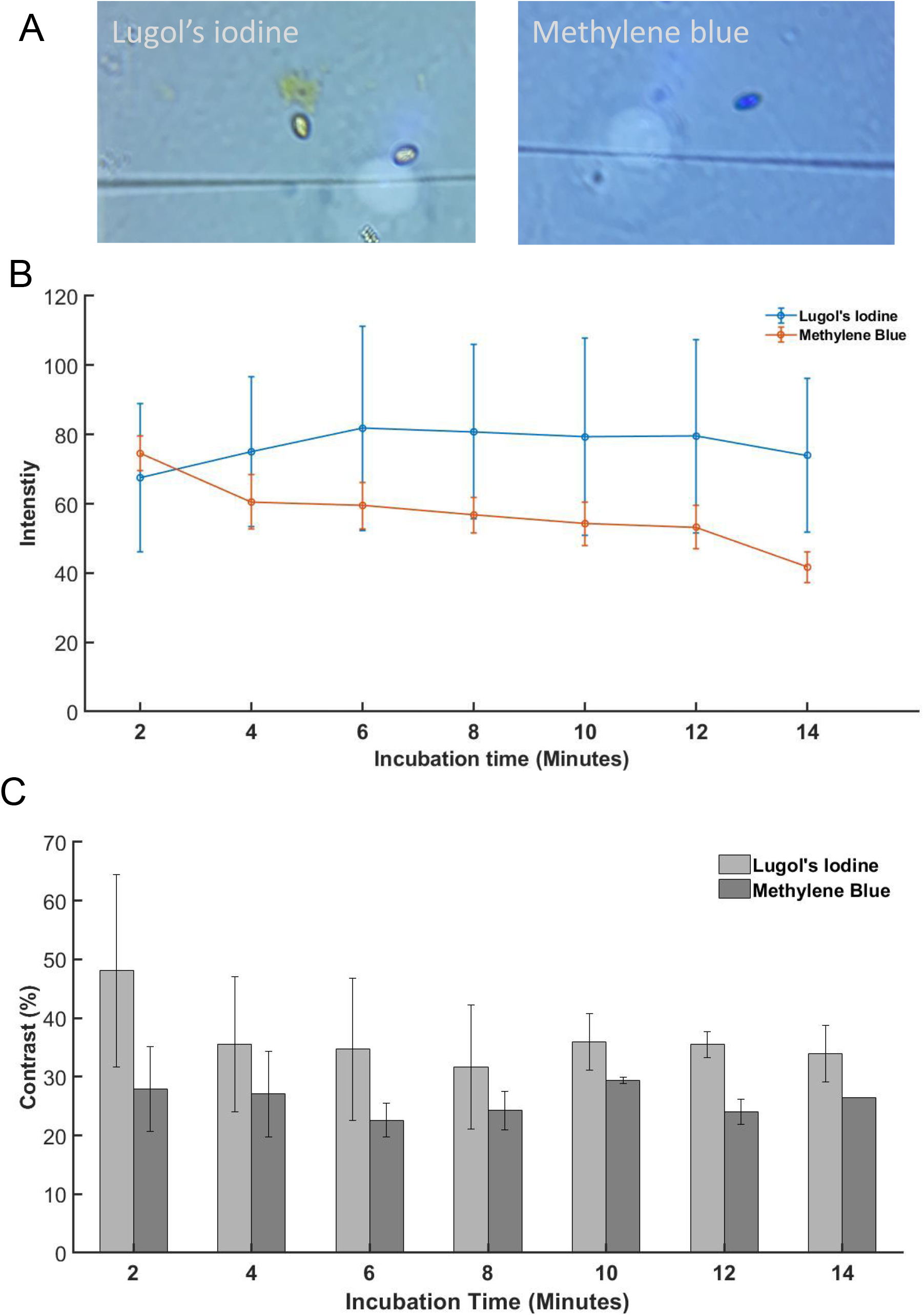
Representative images of Lugol’s iodine and methylene blue staining (A). Both of the images were taken at 10 min of staining. The average intensity of (oo)cysts under white LED light source at different time intervals (B). A plot of contrast versus incubation time for methylene blue (MB) and Lugol’s iodine staining (C).

It is evident that Luglo’s iodine (LI) provided brighter image than Methylene blue (MB). Also, LI staining, intensity increased after 2 minutes of incubation and remained constant for up to 12 minutes. This indicated that LI staining is more stable over time. Based on stability of stain and intensity, we selected LI staining and 6-8 minutes of staining time in the subsequent experiments. The Lugol’s iodine staining also provided higher contrast compared to methylene blue dye staining. Lugol’s iodine initially has contrast of 48±16.4% which decreased to 35.5±11.6% in next 2 minutes and remained constant throughout the time. In case of methylene blue contrast remained constant around 27.7% during the experiment.

### Method validation

Accuracy of the smartphone microscope was evaluated by spike recovery experiments using both vegetable and water samples. In this experiment, known number of (oo)cyst were spiked to the sample and the number of (oo)cyst recovered were counted with the smartphone microscope. We compared the results from smartphone microscope with measurements using commercial bright field and fluorescence microscopes (Table 2).

**Table 2:** Percentage recovery of *Cryptosporidium* and *Giardia* using smartphone, commercial brightfield, and fluorescence microscopes.

| Sample | Recovery (%) |  |  |  |  |  |
| --- | --- | --- | --- | --- | --- | --- |
|  | Smartphone microscope |  | Brightfield microscope |  | Fluorescence microscope |  |
|  | Cryptosporidium | Giardia | Cryptosporidium | Giardia | Cryptosporidium | Giardia |
| Cabbage | 26.8±10.3 | 10.2±4.0 | 40.2±7.1 | 21.6±6.7 | 21.2±3.2 | 18.4±2.6 |
| Carrot | 40.1±8.5 | 14.1±7.3 | 62.7±15.8 | 24.7±11.4 | 22.8±5.5 | 20.1±6.3 |
| Cucumber | 44.4±7.3 | 24.2±12.1 | 51.9±18.3 | 26.7±10.6 | 22.8±3.5 | 19.9±4.1 |
| Radish | 47.6±11.3 | 23.2±13.7 | 60.3±12.3 | 26.7±11.9 | 23.9±5.0 | 20.4±2.9 |
| Tomato | 49.2±10.9 | 17.1±13.9 | 58.3±14.3 | 21.7±9.0% | 32.3±6.6 | 29.2±2.8 |
| Water | 30.2±7.9 | 37.6±2.4 | 35.3±7.0 | 45.6±6.6 | 44.4±8.8 | 55.8±7.4 |

The recovery of *Giardia* ranged from 10.2±4.0% in cabbage to 37.6±2.4% in water and recovery of *Cryptosporidium* ranged from 26.8±10.3% in cabbage to 49.2±10.9% in tomato using smartphone microscope measurement (*see* Table 2). For most of samples, the percentage recovery was found to be marginally higher in bright field microscopy than in smartphone microscopy. The recovery of oocyst was higher than the cyst in all three microscopes with few exceptions. In image plane (in camera), the smartphone microscope has circular field of view having diameter of ~200 μm, whereas the commercial bight filed microscope at 400× has rectangular field of view of ~190 μm×350 μm. The lower percentage recovery in smartphone than in brightfield microscope is mostly likely due to lower field of view which makes (oo)cyst counting difficult. The percentage recovery data reported in this work are comparable to literature studies. Cook *et al*. [30] reported the percentage recovery of *Giardia* and *Cryptosporidium* in spiked lettuce and raspberries of 30.4%. and 44.3%, respectively. In another study, Cook *et al*., [31] developed a method for detection of *Giardia* cysts and *Cryptosporidium* oocysts on lettuces and other salad products. They used texas red dye to stain *Giardia* cyst and *Cryptosporidium* oocyst that yielded recoveries on a variety of commercially available natural foods of 36.5±14.4% and 36.2 ±19.7%, respectively. Similarly, in a study conducted by Amoro *et al*. [25] in 19 salad products, following the same method of Cook *et al*. [30–32], recoveries of the texas red–stained *Cryptosporidium* and *Giardia* were 24.5± 3.5% and 16.7 ±8.1% respectively.

The recovery efficiencies varied in certain percentages among all five different vegetables. The average recovery efficiency of (oo) cysts of *Giardia* by both brightfield microscopy and smartphone microscopy was higher in radish and cucumber. On the other hand, the bright field microscopy showed more *Cryptosporidium* in carrot and radish whereas, it was higher in radish and tomato by smartphone microscopy. The highest recovery efficiency was observed in tomato followed by radish, carrot, cucumber and cabbage for both *Giardia* and *Cryptosporidium* by fluorescence microscopy (Table 2). This difference in recovery efficiencies in various vegetables using the same methodology may be due to the variability of the noncovalent interactions between (oo) cyst surfaces and surfaces of various vegetables we tested. It is also important to note that the extraction methods that have been proven suitable for one specific food matrix could be unsuitable for the others [31–34]. For example, the glycine wash buffer had satisfactorily recovered both *Giardia* and *Cryptosporidium* in lettuce and raspberries [13, 31–33]. On the other hand, a prolonged, vigorous washing of *Spinacia oleracea* [35] and apples [36] in 1 M glycine (pH 5.5) elution buffer was not able to remove all of the *Cryptosporidium* oocysts from their matrix[35, 36].

For comparison, we also performed spike recovery experiments with a commercial florescent microscope. The highest recovery was observed in tomato samples for *Cryptosporidium* (32.3±6.5%, n =15) and for *Giardia* (29.3 ± 2.8%, n =15)which was followed by radish samples with (23.9 ±5.4%, n=15) for *Cryptosporidium* and (20.4±2.9%, n=15) for *Giardia* (Table 2). The recovery efficiency was consistently lowest for all the samples with the fluorescence microscopy except for the tomatoes and water samples, in comparison to the smartphone and bright field microscopy. The recovery using fluorescence microscope, in which the (oo)cysts were tagged with fluorescent dye tagged antibody, was found to be lower than in remaining two microscopes, except in water samples. Fluorescence microscopy is dependent on binding of the fluorescence tagged antibodies to the antigen surface, which can be hindered/altered by the impurities present in the solution like the vegetable debris in our case. In some cases, fluorescent antibody could bind to the impurities and show the false positive. This could be an explanation for the higher fluorescence in the sample purified from the tomatoes. In addition of hindering binding of antibodies to oo(cysts), the larger particles can deposit (oo)cysts underneath so that they are no more accessible for antibodies. So, lower detection in case of radish, cabbage, cucumber and carrot could be due to decreased binding affinity of the antibody to the (oo)cysts or hiding of (oo) cysts underneath the larger vegetable debris [37].

Table 2 also lists the recovery efficiencies of the spiked water samples, detected by the smartphone microscope with a commercial bright filed microscope and fluorescence microscope. The recovery of both *Giardia* cysts (55.9±7.4%) and *Cryptosporidium* oocysts (44.4±8.8%) were higher in fluorescence microscopes compared to both smartphone and bright field microscopes. For smartphone microscope, 37.6±2.4% cysts and 30.2%±7.9% oocysts were observed whereas it was 45.6±6.6% cysts and 35.3±7.0% oocysts in bright field microscopy.

Previous studies have reported similar percentage recovery data. Le Chevallier et al. used the immunofluorescence microscopic method and reported an average recovery efficiency of 68.6% for *Giardia* cysts and 25.3% for *Cryptosporidium* oocysts in seeded tap water [38]. In another study Le Chvallier et al. showed a recovery of 96% and 77% for cysts and oocysts respectively, with the Percoll sucrose density gradient at a specific gravity ≥ 1.10 [39]. Koompapong et al. using a similar methodology reported a recovery of oocysts (75%) in water samples [40]. In contrast, Machado et al. found a significantly small recovery of 5.3%, who analyzed the sediment of water samples using Kinyoun and Koster histochemical staining techniques [41]. They didn’t use any chemical precipitant for the flocculation of oocysts before purification steps. Karanis et al (2001) compared different flocculants and concluded that using ferric sulfate yield a higher recovery (61.5%) of *C. parvum* oocysts from tap water with a very low impact on the viability of oocysts [27]. Also, no detergent solutions were included in the study that helped to set the oocysts free from the sediments [41]. In a study made by Hsu et al. standard Envirochek capsule filtration followed by immunomagnetic separation, the standard purification procedure in Environmental Protection Agency Method 1623, was used. In their study, the recovery efficiencies were higher for *Giardia* (48.0%) than for *Cryptosporidium* (32%) [42]. These data are very similar to our percentage recovery data in water samples.

We also estimated method detection limit (LOD) of smartphone microscope method. The LOD varied with type of sample. LOD of *Giardia* ranged from 24 cyst/100 g for cucumber to 73 cyst/100 g for cabbage (tomato = 38, carrot = 40 and cucumber = 23 cyst/100 g). Similarly, the LOD for *Cryptosporidium* ranged from 11 oocyst/100 g for radish to 25 oocyst/100 g for cabbage (tomato = 12, carrot = 12 and cucumber = 23 oocyst/100 g). In general, the LOD of *Cryptosporidium* was lower than that of *Giardia*.

### Prevalence of (oo)cysts in vegetable and water samples

After developing the smartphone microscopic system for (oo)cyst detection, we screened (oo)cysts contamination in five different types of vegetable samples (n = 196) purchased from local market in Kathmandu, Nepal and river water samples (n=18). The samples were processed as described in method section.

The result is shown in figure 4. Out of the total samples, 31.1% vegetable samples were positive for cyst and 35.2% samples were positive for oocyst contamination when detected using smartphone microscope. The prevalence of oocysts was highest in spinach samples and lowest in carrot samples (*see* Figure 4). The same samples were also tested using brightfield microscope and fluorescence microscope and compared with smartphone microscope. The comparison is provided in Table 3. Cysts were detected in 40.3% samples oocysts were detected in 41.8% vegetable samples using brightfield microscope. Further, out of 196 samples, randomly selected 58 (30%) samples were screened using fluorescence microscope. The fluorescence measurement showed that 27(46.5%) samples were contaminated with cyst and 26 (44.8%) samples were contaminated with oocyst. The shape and surface of vegetables might play a role leading to the contamination. The (oo)cysts can easily attach to the uneven or curly surfaces of spinach and cabbage either in the farm or when washed with polluted water. On the other hand, vegetables with smooth surfaces such as radishes and carrots had a low number of (oo)cysts in the present study as its smooth surface reduces the attachment of the protozoans [44].

**Figure 4:**
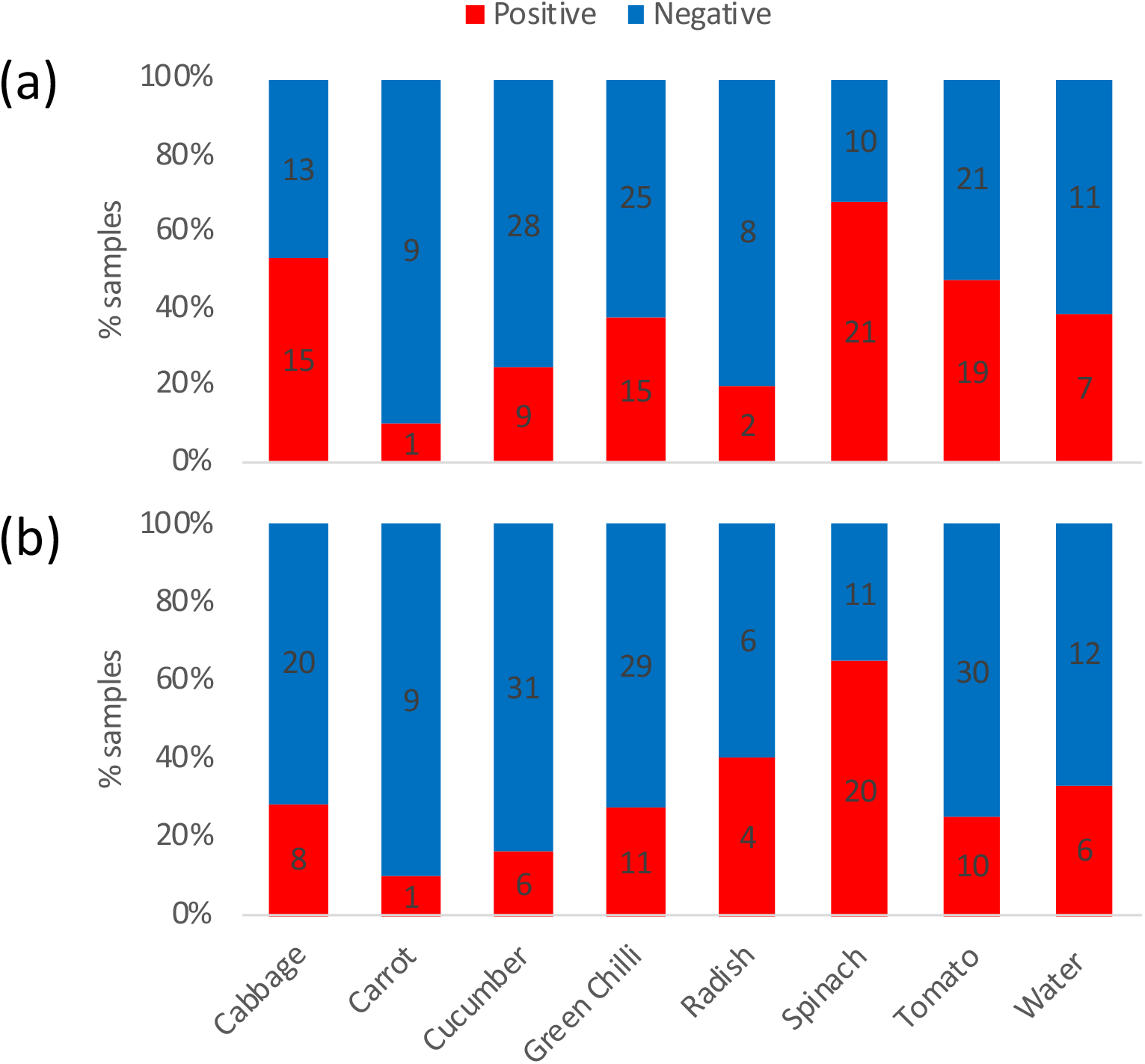
Percentage of presence or absence of cyst in each sample types using smartphone microscope: a) Cryptosporidium and b) Giardia. The numbers in each bar graph represent the number of samples.

According to a survey conducted by Maikai *et al*. 40% of Spinach, 32% of tomato, 24% of carrot and 16% of cabbage were contaminated with *Cryptosporidium* oocysts using microscopy [45]. A study by Kaudah *et al*. reported that 50 % tomatoes, 43.1% cabbage and 26.4% of carrot were tested positive for different protozoans among which 11% were *Cryptosporidium*. These samples were stained with Lugol’s iodine and observed under light and fluorescence microscope [46]. In contrast Utaaker *et al*. reported that only 14% (8/56) tomatoes and 9% (4/47) cabbages were contaminated with either *Giardia* cysts or *Cryptosporidium* oocysts [22]. In case of root vegetables such as carrot, the current study has a very low record (10%) for both *Giardia* and *Cryptosporidium*. Similar results were also reported in other studies like 14% positive cases in India [22] and 6.4% in Southern Ethiopia [47] for *Giardia*. A slightly higher positive cases was observed in Egypt (43.3%) [44] and Korea (33.3%).

We tested eighteen surface water samples collected from 3 different sites of the Bishnumati river, Kathmandu, Nepal in two different field campaigns. The samples were flocculated and purified with sucrose density gradient and examined by smartphone, commercial bright field, and fluorescence microscopies. A total of 33.3% (6 out of 18 samples) were tested positive for *Giardia* and seven samples (38.9%) were tested positive for *Cryptosporidium* by smartphone microscope (figure 4). When compared to other microscopes, in general, higher number samples were tested positive for (oo)cysts by the smartphone microscopy. Brightfield microscope confirmed 22.2% (4 out of 18) positive results for *Giardia* and 33.3% (6 out of 18) positive results for *Cryptosporidium*. Similarly, four (22.2%) were tested positive by for (oo)cysts using fluorescence microscopy (*see* Table 3).

**Table 3:** Prevalence of (oo)cyst contamination in vegetable and water samples measured by three different microscopic methods.

| *Sample type* |  | *Number of samples* |  |  |  |  |  |
| --- | --- | --- | --- | --- | --- | --- | --- |
|  |  | *Smartphone*<br>*microscope* |  | *Brightfield*<br>*microscope* |  | *Fluorescence*<br>*microscope* |  |
|  |  | *Crypto* | *Giardia* | *Crypto* | *Giardia* | *Crypto* | *Giardia* |
| Cabbage | *positive* | 15 | 8 | 9 | 11 | 6 | 4 |
|  | *negative* | 13 | 20 | 19 | 17 | 9 | 5 |
| Carrot | *positive* | 1 | 1 | 1 | 1 | 1 | 1 |
|  | *negative* | 9 | 9 | 9 | 9 | 4 | 0 |
| Cucumber | *positive* | 9 | 6 | 6 | 9 | 3 | 5 |
|  | *negative* | 28 | 31 | 31 | 28 | 8 | 4 |
| Green<br>Chilli | *positive* | 15 | 11 | 15 | 14 | 6 | 2 |
|  | *negative* | 25 | 29 | 25 | 26 | 12 | 9 |
| Radish | *positive* | 2 | 4 | 2 | 3 | 0 | 3 |
|  | *negative* | 8 | 6 | 8 | 7 | 0 | 0 |
| Spinach | *positive* | 21 | 20 | 20 | 21 | 6 | 6 |
|  | *negative* | 10 | 11 | 11 | 10 | 8 | 10 |
| Tomato | *positive* | 19 | 10 | 15 | 20 | 3 | 4 |
|  | *negative* | 21 | 30 | 25 | 20 | 8 | 5 |
| Water | *positive* | 7 | 6 | 6 | 4 | 4 | 4 |
|  | *negative* | 11 | 12 | 12 | 14 | 14 | 14 |

We estimated the (oo)cysts per unit of sample. The highest concentration of the (oo)cysts for both *Giardia* and *Cryptosporidium* were detected in cabbages (n= 27) with the concentration of 442 cysts and 225 oocysts/kg. The lowest concentration of 35 cysts/kg and 16 oocysts/kg was found in radish. Tomatoes (n= 40), carrots (n=10), and cucumbers (n= 37) were found to be contaminated with 129 cysts/kg, 166 cysts/kg, and, 77 cysts/kg, respectively. Similarly, 76 oocysts/kg, 47 oocysts/kg, 185 oocysts/kg, and were found in tomato (n=40), carrots (n= 10), and cucumber (n=37), samples. The infectious dose for cryptosporidiosis and giardiasis is as low as 10–30 viable (oo)cyst [48]. Assuming around 200 g of poorly washed raw vegetable is consumed per day, there is still high chance that most of the samples tested were infectious if ingested.

## 4. Conclusions

We designed a smartphone microscope and optimized its various optical parameters. The field of view increases with the diameter of sapphire ball lens but the magnification follows the opposite trend; in agreement with theory. We found that microscope having ball lens of 1 mm diameter along with Lugol’s iodine staining and commercially available white LED illumination can simultaneously determine (oo)cyst of *Cryptosporidium* and *Giardia* in vegetable samples. The spiking recovery experiment on the different vegetable and water samples showed that the % recovery is comparable to the commercial bright field microscope and better than fluorescence microscopic measurement. We found that % recovery varied with the nature of sample and recovery for *Cryptosporidium* oocyst is better than *Giardia* cyst. This observation is consistent with the literature studies. Out of the 196 vegetable samples 31.1% vegetable samples were positive for cyst and 35.2% samples were positive for oocyst contamination when examined by smartphone microscope. This study shows that the smartphone based microscopic assay can a low-cost alternative for pre-screening of (oo)cyst of *Cryptosporodium* and *Giardia* in resource limited settings. Our future work involves the development of an automated smartphone program that could take image, process the image to identify and count the (oo)cysts, and provide report to the user. This automated system may minimize error and shorten the analysis time.

## Acknowledgments

This work was supported by NAS and USAID through Partnerships for Enhanced Engagement in Research (PEER) (AID-OAA-A-11-00012). The opinions, findings, conclusions, or recommendations expressed in this article are those of the authors alone, and do not necessarily reflect the views of USAID or NAS.

